# M-CSF-stimulated myeloid cells can convert into epithelial cells to participate in re-epithelialization and hair follicle regeneration during skin wound healing

**DOI:** 10.1101/2021.12.17.473218

**Authors:** Yunyuan Li, Hatem Nojeidi, Ruhangiz T. Kilani, Aziz Ghahary

**Affiliations:** Department of Surgery / Plastic Surgery, University of British Columbia, Vancouver, BC, V5Z 1M9 Canada

**Author notes:** **Correspondence and requests of materials should be addressed to**: Aziz Ghahary, PhD, Professor, 4520 -ICORD, 818 10^th^ Avenue, Vancouver, BC, V5Z 1M9 Canada.

## Abstract

Skin wound healing is a complex process which requires the interaction of many cell types and mediators in a highly sophisticated temporal sequence. Myeloid cells compose a significant proportion of the inflammatory cells recruited to a wound site and play important roles in clearance of damaged tissue and microorganisms. Myeloid cells have also been suggested to convert into fibro-blast-like cells and endothelial cells in participation of wound healing process. However, whether myeloid cells in wound skin can convert into epithelial cells and contribute to re-epithelialization and skin appendage regeneration is still unclear. In this study, we performed double immunofluorescent staining with antibodies for hematopoietic cells and keratinocytes as well as cell tracing technique to investigate hematopoietic cell conversion. The result show that during the healing process, some of the CD45-positive hematopoietic cells are overlapped with keratin 14, the markers of keratinocytes. Further, CD11b-positive myeloid cells seem the origin of converted epithelial cells. To confirm these results, we culture CD11b-posiitve myeloid cells from mouse splenocytes in a medium containing macrophage colony-stimulating factor (M-CSF) and dermal injection of these cells into the healthy skin when punch biopsy is created in mouse skin. Tracing injected labeled splenocyte-derived myeloid cells in skin, we confirm that myeloid cells able to convert into keratinocytes in repaired skin. Furthermore, our results from *in vivo* experiments provide new information on contribution of myeloid cells in hair follicle regeneration. In conclusion, this work highlights the myeloid cell contributions in wound repair and hair follicle regeneration in mice through conversion of M-CSF-stimulated CD11b+ myeloid cells into epithelial cells.

## Introduction

Dermal wound healing is a dynamic process involving the coordination of epidermal keratinocytes, dermal fibroblasts, endothelial cells and infiltrated hematopoietic cells. After injury, it is thought that fibroblasts residing in the reticular dermis of wound edge contribute to the dermis replacement while basal keratinocytes and epithelial stem cells accomplish the re-epithelialization through cell migration and proliferation [1]. It may be true for the healing of a small skin wound. However, a large full-thickness wound is unlikely to heal solely just through migration and proliferation of skin cells resided at the wound edge. Last few years, we have proposed and begun to investigate whether a subset of infiltrated hematopoietic cells have a capacity to be converted into skin cells or stem cells and thereby contributing to skin wound healing. In fact, in previous publication, our result showed that skin injection of M-CSF-cultured adherent hematopoietic cells can accelerate skin wound healing [2]. Other studies demonstrated that in addition to fibroblasts migrated from the wound edges, monocyte-derived fibrocytes from circulation [3] and residing macrophages converted into fibroblast-like cells also contribute to dermal tissue repair [4].

During the healing process in different tissues, it has been noticed that hematopoietic stem cells or hematopoietic cells have the capacity to be transdifferentiated into tissue specific cells such as neuronal cells, endothelium, skeletal muscle, hepatocytes and epithelial cells [5–9]. Although the capacity of hematopoietic stem cell or hematopoietic cell transdifferentiation has been questioned by some studies indicating cell fusion of hematopoietic cells with other cells rather than transdifferentiation *in vivo*, and mimic appearance of transdifferentiation from grafted hematopoietic cells after transplantation [10–12], new evidence from recently study by using cell tracing technique and single cell sequencing strongly support that hematopoietic cells specifically myeloid cells can convert to solid organ specific cells. For example, myeloid cells in wound bet have been demonstrated to convert to fibroblast-like cells [4] while csf1r-expressed erythro-myeloid progenitors can convert to endothelial cells in blood vessels during embryo development and in adult [13].

To further assess whether infiltrated hematopoietic cells in wound bed can convert to epithelial cells, contributing to epithelialization and skin appendage regeneration, here, we used antibodies of hematopoietic cells and keratinocytes for double immunofluorescent staining as well as cell tracing approach to address this question. The result showed that infiltrated myeloid cells could not only convert to keratinocytes for repairing damaged skin but also contribute to hair follicle regeneration in a mouse skin wound healing model.

## Materials and Methods

### Mice

Balb/C and C57BL/6 were purchased from The Jackson Laboratory (Bar Harbor, Marine). All experimental procedures were approved by the University of British Columbia Animal Care Committee. The methods were carried out in accordance with the approved guidelines.

### Skin wound model

Two 8 mm punch biopsy excisional wounds were created on the dorsal skin of mouse after mouse was anesthetized by low-dose isoflurane inhalation. To prevent contraction of wounds, silicon splints were used to allow wounds to heal through granulation and re-epithelialization.

### Immunofluorescent staining

Skin samples were fixed with 10% formalin. About 5 μm thick sections were cut and used for immunofluorescent staining. Following deparaffinization and antigen retrieval, skin sections were incubated overnight with primary antibody after blocking non-specific binding with blocking solution containing 5% bovine serum albumin for one hour. For cell staining, cells were fixed with 10% formalin, blocked with blocking solution and incubated with primary antibody overnight without performing antigen retrieval. After washing three times with PBS or PBS-T (0.1% Triton X-100 in PBS) at room temperature, sections or cells were incubated with secondary, fluorescein-conjugated antibodies for another one hour. After washing three times, cells (or sections) were stained with 1.5% Vectashield mounting medium containing DAPI (Vector Laboratories). Images were captured using a Zeiss Axioplan 2 fluorescence microscope and AxioVision image analysis software. The primary and secondary antibodies used in this study were listed in Supplemental Table S1.

### Isolating and culturing cells from mouse skin

Skin of mouse was harvested and washed three times with phosphate buffer saline (PBS) containing three-fold antibiotic-antimycotic (Invitrogen Life Technologies, Carlsbad, CA). Skin was minced into small pieces (about 2-3 mm × 2-3 mm) and digested by incubation with 1 mg/mL collagenase V (Sigma) in PBS for one hour with shaking (250 rpm). The collagenase was then neutralized by adding an equal volume of Dulbecco’s Modified Eagle Medium (DMEM, Invitrogen Life Technologies) with 10% fetal bovine serum (FBS, Invitrogen Life Technologies). Cells were filtered through a 70 μm cell strainer and washed twice with DMEM containing 10% FBS. Cells were cultured in DMEM with 2% FBS for further experiments.

### Culturing myeloid cells

Splenocytes were isolated from Balb/C or C57BL/6-Tg (UBC-GFP) mice. GFP mice were kindly provided by Mr. Darrel Trendall at Jack Bell research Centre, University of British Columbia. Splenocyte-derived myeloid cells were generated by culture in a medium containing 5 ng/ml M-CSF. Cells in passage two were used for cell injection to the mice.

### Labeling cells *in vitro* by Dil prior to injection

Splenocyte-derived, M-CSF-cultured myeloid cells at passage 1-2 were used for this experiment. Cells were detached by trypsin-EDTA, spun down and then re-suspended in 1 mL PBS containing 20 μg/mL Dil and incubated for 15 minutes at room temperature. Unbound Dil was then saturated with 1 mL of FBS and removed by centrifugation and washing three-times with cell culture medium. Dil-labeled cells in PBS were ready for injection.

### Dermal injection of myeloid cells to the healthy skin of wounded mice

When excisional wounds were created by punch biopsy, one million Dil-labeled or GFP expressing myeloid cells in 100 μL PBS were dermally injected simultaneously at one spot on the dorsal healthy skin, one cm far from the edge of wounds.

### Statistical analysis

All data were presented as the mean ± standard deviation. Statistical analyses were performed with GraphPad inStat software. *P* values were calculated using two-tailed unpaired student’s t test. A p value of <0.05 was considered as statistically significant.

## Results

### Distributions of hematopoietic cells in healed skin wounds

It is well known that hematopoietic cells such as neutrophils, lymphocytes and monocytes can infiltrate to the wound sites during the healing course after skin injury [14]. We first investigated whether hematopoietic cells are still present in the wounded skin after wounds are completely healed. To do this, we created dermal wounds by an eight mm size punch biopsy at dorsal skin of mice (Total 16 wounds were created from 8 Balb/C mice), used silicon splint to prevent wound contraction, and collected skin samples at day 14 (wounds were closed around day 10-12). As shown in Figure 1A, using an antibody for hematopoietic cell marker, CD45 to perform immunofluorescent staining in wounded skin of mice, a lot of CD45-positive cells were revealed at not only dermis but also epidermis of 3 of 16 complete epithelialized wounds. To compare the different of CD45-positive cell number in normal skin and wounded skin of mice. Five normal skin and five wounded skin were performed an immunofluorescent staining with CD45 and the ratio of CD45-positive cell number and the total cell number (DAPI stain, blue) was calculated. A markedly higher number of CD45-positive cells in wounded skin was found as compared to that in uninjured normal skin (55.6 ± 12.3 % vs 4.5 ± 1.3 %; wounded skin vs Normal skin, n=5) (Fig. 1B).

**Figure 1.**
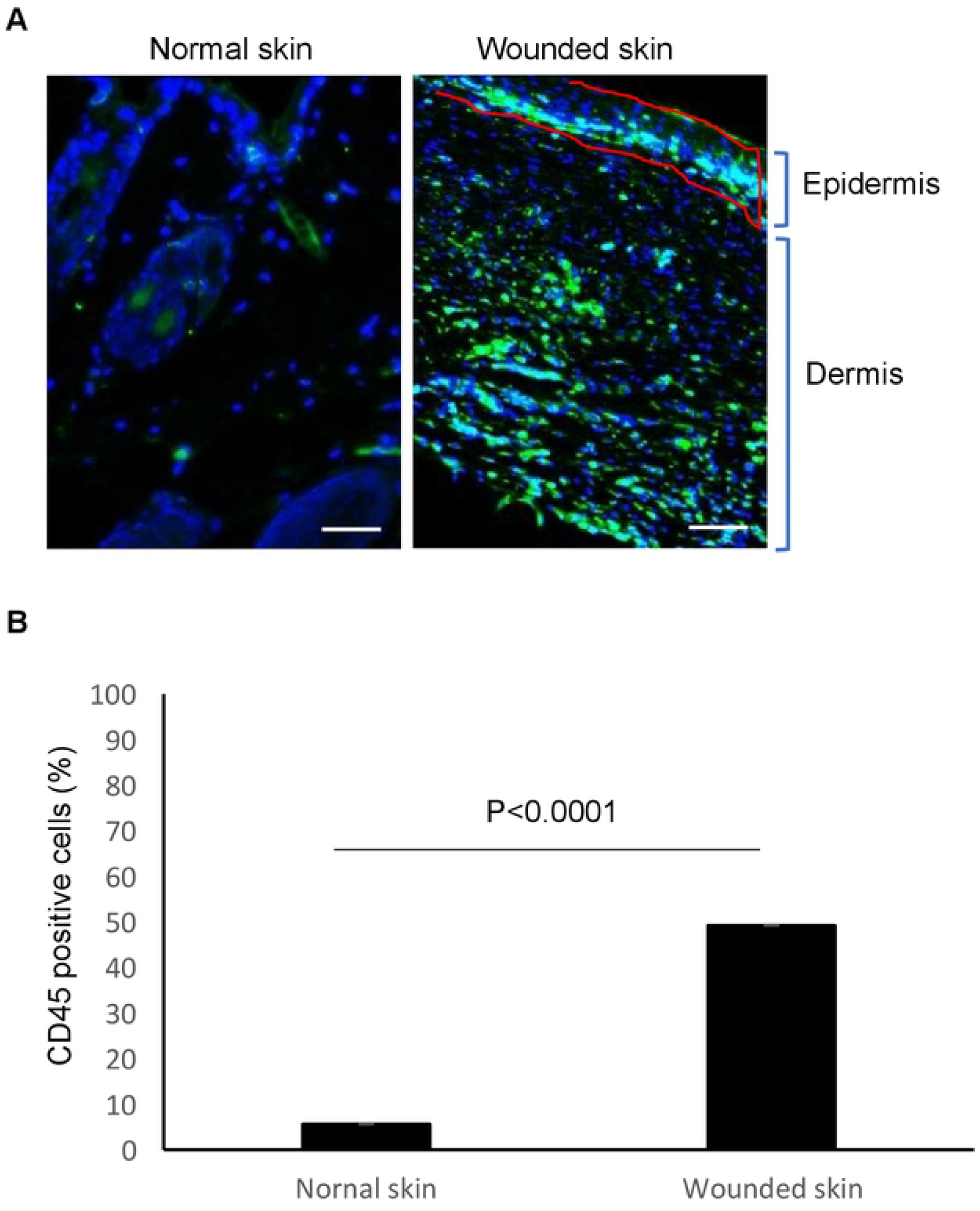
CD45-positive hematopoietic cells in skin wounds of mice. (**A**) Immunofluorescence stain in normal and wounded skin of mice with CD45 antibody. Balb/C mice with 8 mm punch biopsy excisional wounds for 14 days were euthanized and wounded skin and normal dorsal skin were harvested, fixed and paraffinized. Skin sections were prepared for immunofluorescent staining with CD45 antibody (green). DAPI (blue) was used as a nuclear counterstain. Scale bars in all images was 50 μm. A representative of wounded skin of 16 wounds from 8 mice was shown. (**B**) The statistical analysis of CD45 positive cell percentage in skin sections is depicted. Ten HPF of five normal and five wounded skin were counted (n=5).

### Conversion of hematopoietic cells to keratinocytes in repaired epidermis of skin

As shown in Figure 1A, some of CD45 positive hematopoietic cells were distributed at epithelialized epidermis. Here, we further investigated whether infiltrated hematopoietic cells also able to convert to keratinocytes or keratinocyte-like cells. To address this question, tissue sections from healed skin wounds and normal skin of mice were performed a double immunofluorescent staining with antibodies for CD45 and keratinocyte marker, keratin 14 (K14). The result showed that some double positive cells were found at the basal layer of re-epithelialization epidermis while none of double positive cells were found in uninjured normal skin (Figure 2A). To confirm this result, we further isolated skin cells from wounded skin by collagenase digestion, cultured for 48 hours and performed a double immunofluorescent staining with antibodies for CD45 and K14. As shown in Figure 2B, in consistent with the result shown in Figure 2A, some of the K14 positive keratinocytes were also positive for CD45, suggesting that some of CD45-positive hematopoietic cells really convert to epithelial cells to participate in epithelialization after injury.

**Figure 2.**
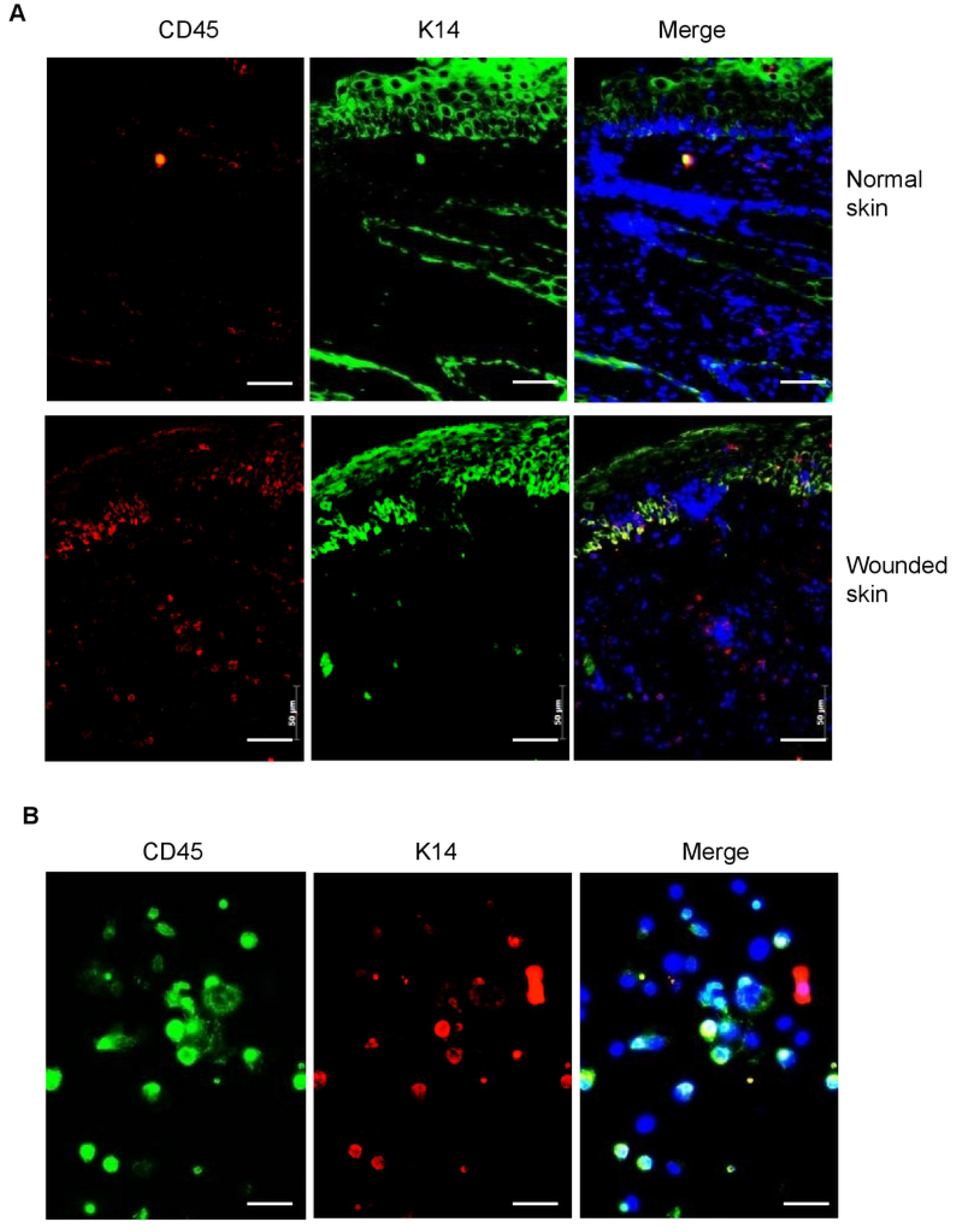
Conversion of CD45-positive hematopoietic cells to keratinocytes in re-epithelialization skin of mice. (**A**) Double immunofluorescent staining with keratin-14 (K14) and CD45 antibodies in normal skin (Top panels) and wounded skin (Bottom panels) of Balb/C mice. Wounded skin was taken from mice with 8 mm punch biopsy of excisional wound after wounding for 14 days. (**B**) Double immunofluorescent staining with K14 and CD45 antibodies in cultured skin cells isolated from wound edge of Balb/C mouse skin. Skin cells were isolated from the edge of wounded skin of mice by collagenase digestion. Cells were cultured for 48 hours before fixation and performing an immunofluorescent staining. DAPI (blue) was used as a nuclear counterstain. Scale bars in all images was 50 μm.

### Myeloid cells are the cellular origin of hematopoietic cell-converting to keratinocytes in wounded skin of mice

As myeloid cells have been suggested to be the cellular origin of converted fibrocytes [3], fibroblast-like cells [4] and endothelial cells [13], here, we examined whether these skin keratinocytes converted from infiltrated CD45-positive hematopoietic cells are originated from myeloid cells. By using double immunofluorescent staining with antibodies CD45 and myeloid cell marker CD11b to detect wounded skin of mice, we found the most of CD45-positive cells distributed in repaired epidermis were also positive for CD11b (Fig.3A). We then performed a double immunofluorescent staining with CD11b and K14 antibodies in wounded skin of mice. The result showed that some of CD11b-posiitve cells located near basal membrane layer of epidermis were also stained K14 (Fig. 3B). Similar result was obtained in cultured skin cells from wounded skin of mice (Fig. 3C). These results suggest that CD11b-positive myeloid cells are the origin of keratinocytes converted from hematopoietic cells.

**Figure 3.**
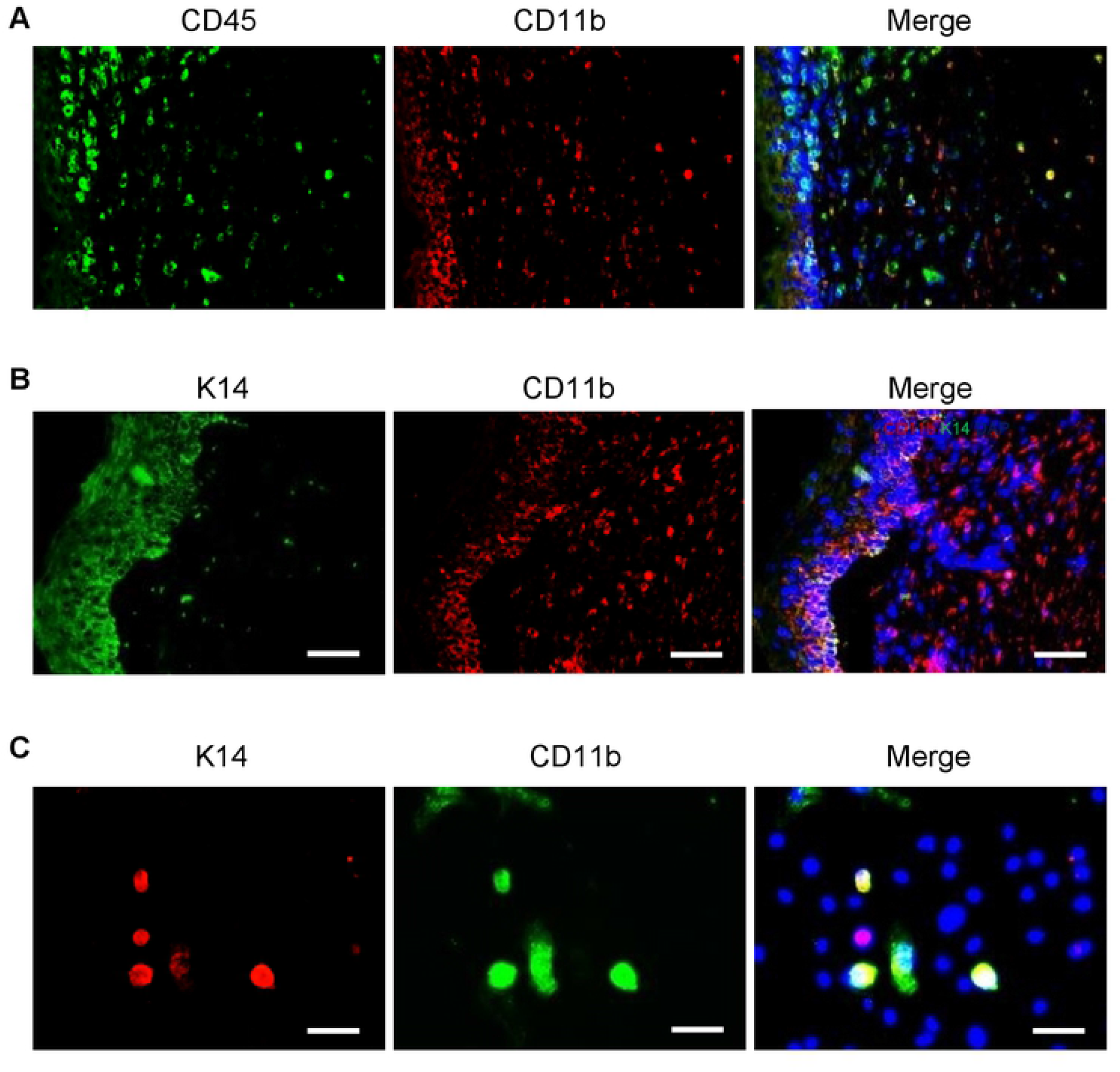
Myeloid cells are the cellular origin of hematopoietic cell-converting keratinocytes. (**A**) Double immunofluorescent staining with antibodies of CD45 and myeloid cell marker CD11b in wounded skin of mice. (**B**) Double immunofluorescent staining with antibodies of CD11b and K14 in wounded skin of mice. (**C**) Double immunofluorescent staining with antibodies of CD11b and K14 in cultured skin cells. Skin cells were isolated by collagenase from wounded edge skin of Balb/C mice after injury for 14 days and cultured for 48 hrs in vitro before fixation and stained. DAPI (blue) was used as a nuclear counterstain. Scale bars in all images was 50 μm.

To provide evidence that M-CSF-cultured myeloid cells have the capacity to convert to keratinocytes *in vivo*, splenocytes from normal Balb/C mice were isolated, cultured in M-CSF medium and attached cells were labeled with 1,1’-Dioctadecyl-3,3,3’,3’-Tetramethylindocarbocyanine Perchlorate (Dil). We also used a similar approach to culture myeloid cells from GFP mice but without Dil labelling. More than 94% cells cultured in this condition were CD11b positive while none of adherent cells were positive for Pro-col (fibroblasts) or K14 (keratinocytes) (Figure S1). One million of either Dil-labeled or GFP-expressing syngeneic myeloid cells were then intradermally injected to the normal skin one cm far from the edge of wounds which were generated by 8 mm punch biopsy on the dorsal skin of either the Balb/C or C57BL/6 mice, respectively (Figure 4A). The same strain of mice without cell injection were also wounded as controls. Skin samples from healed wounds were collected after 14 days, skin sections were deparaffinized (for skin of all mice) and further performed immunofluorescent staining with anti-GFP antibody (only for skin of C57/BL mice). As shown in Figure 4B, Dil-labeled myeloid cells (red color under the fluorescent microscope) migrated to the dermis and epidermis of epithelialized wounds but not in mice without cell injection. Similar result was obtained in C57BL/6 mice received GFP-expressing myeloid cells (Figure 4C).

**Figure 4.**
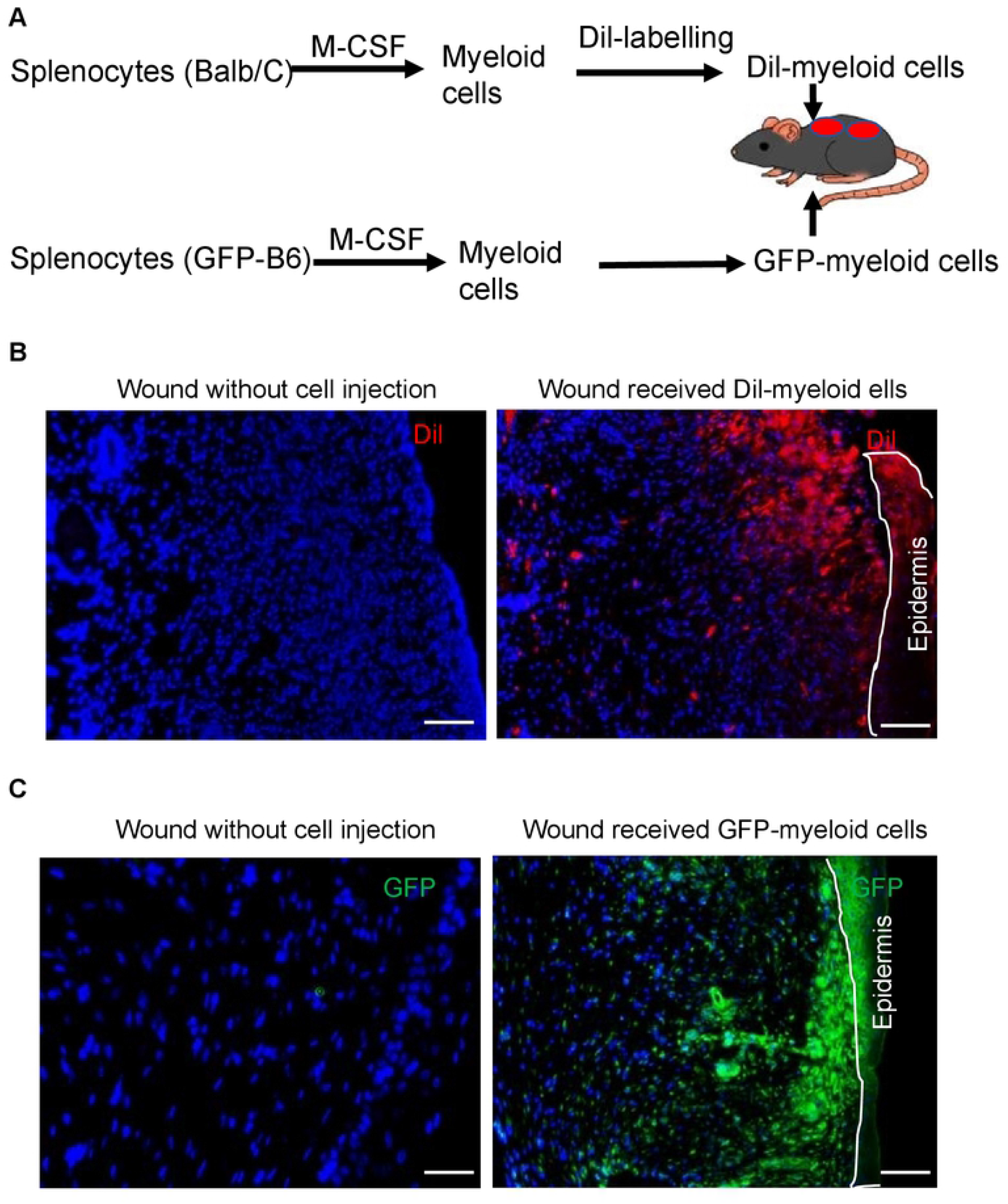
Distribution of injected splenocyte-derived myeloid cells in healed mouse skin. (**A**) Schematic figure of Dil-labeled or GFP-expressing mouse myeloid cell lineage tracing experiments. Splenocytes were isolated from the spleen of Balb/C or C57BL/6-Tg (UBC-GFP) mice and cultured in a medium containing 5 ng/ml of M-CSF to generate myeloid cells. One million labeled cells were dermally injected into the healthy skin at one spot where was one cm far from the edge of wounds at the same time of skin injury. (**B**) Dil-labeled cells in wounded skin of mice received nothing (Left) or one million Dil-labeled splenocyte-derived myeloid cells (Right). (**C**) GFP-positive cells in wounded skin of mice received nothing (Left) or one million GFP-expressing splenocyte-derived myeloid cells (Right). Skins were stained with GFP antibody (green). DAPI (blue) was used as a nuclear counterstain. Scale bars in all images were 100 μm.

To further confirm that injected myeloid cells really converted to skin keratinocytes, we performed double immunofluorescent staining with GFP and K14 antibodies in healed wounds of C57/BL mice which have received GFP-expressing splenocyte-derived myeloid cells. As shown in Figure 5A, result revealed that some of K14 positive keratinocytes were also GFP-positive, suggesting these skin cells were from myeloid cell conversion. Furthermore, the similar result was obtained in cultured skin cells from wounded skin of mice which received GFP-expressed, M-CSF-cultured myeloid cells for 14 days (Figure 5B).

**Figure 5.**
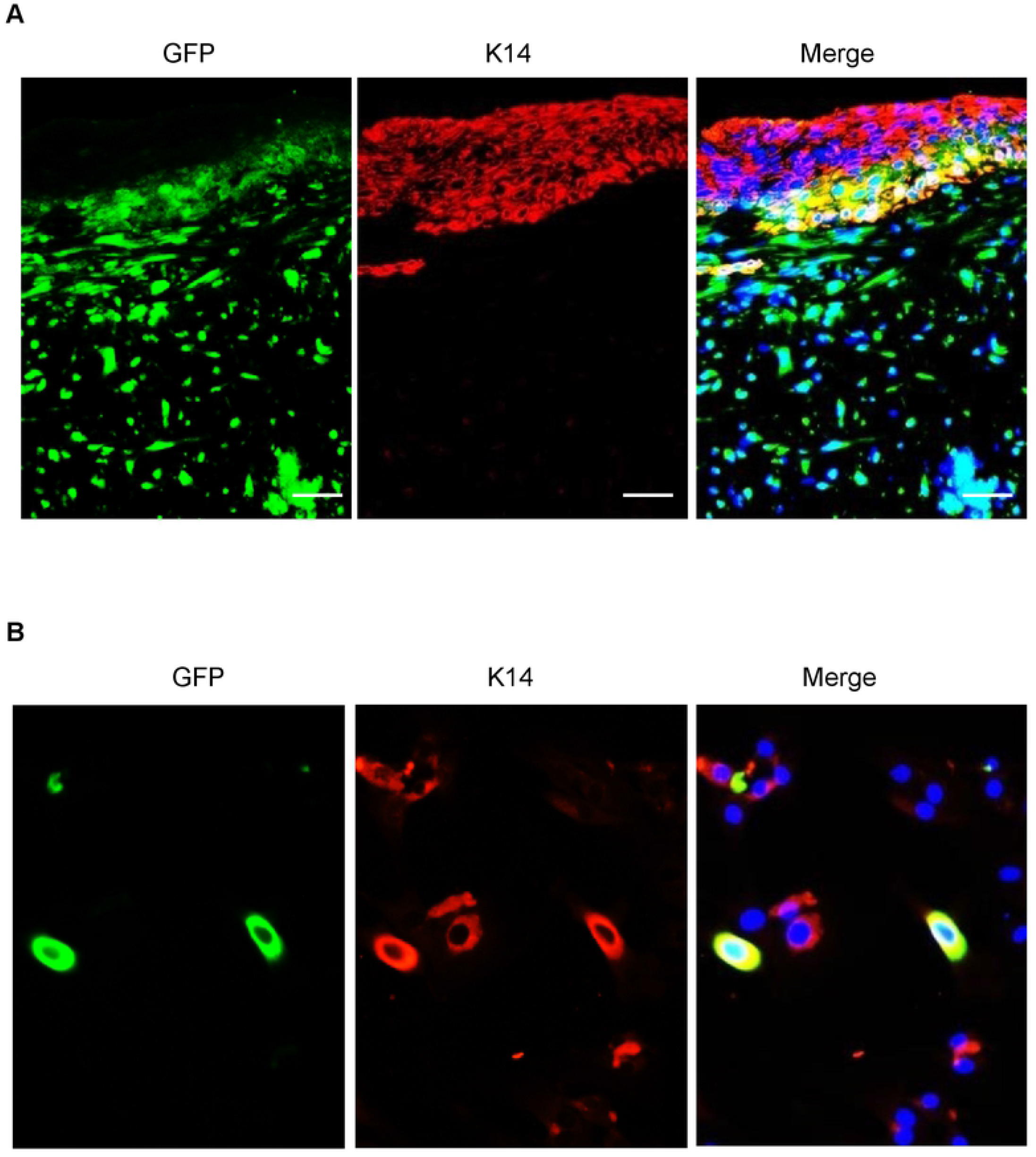
Conversion of GFP-expressing myeloid cells to keratinocytes in re-epithelialization skin. (**A**) Healed skin from wounds of C57BL/6 mice received one million GFP-expressing splenocyte-derived myeloid cells was used for double immunofluorescent staining with antibodies of GFP (green) and K14 (red). (**B**) Double immunofluorescent staining with K14 (red) and GFP (green) antibodies in cultured skin cells. Skin cells were isolated from wound edge skin of mice received GFP-expressing myeloid cells and injury for 14 days. Cells were cultured for 48 hrs before fixation and performing staining. DAPI (blue) was used as a nuclear counterstain. Scale bars in all images was 50 μm.

### Myeloid cells contribute to hair follicle regeneration in re-epithelialization mouse skin

Previous studies have demonstrated that skin injury could induce hair follicle regeneration in some types of adult mice when the size of skin wound is big enough (more than 1 cm) [16–18]. We have also noticed that many CD45-posiitve cells were found in the hair follicles at the edge of wounds and some follicle-like structures were formed in healed skin (Supplement Figure S2). Since we have demonstrated that myeloid cells able to convert into keratinocytes, here, we were interested to further investigate whether myeloid cells participate in regeneration of hair follicles in adult C57BL/6 mice after skin injury. To address this question, two wounds with 8 mm diameter were created by punch biopsy on dorsal skin of each C57BL/6 mouse (total of 5 mice). At the same time of wounding, those mice were dermally injected one million GFP-expressing splenocyte-derived myeloid cells cultured by M-CSF medium at one cm far from the edge of wounds. A silicone wound splint was used on the top of each wound to prevent contraction during the healing. After 4 weeks (a time line for hair follicle regeneration, indicated by previous studies [16–18]), wounded skin was collected for histological examination. We also collected wounded skin to isolate skin cells and hair follicles by collagenase digestion. As shown in Figure 6A, new hair follicle-like structures with green fluorescence in wounded skin were revealed but not in the normal skin from the same mice after 4 weeks post injury. This result was further supported by H & E staining showing new hair follicle-like structures formed in healed wounds (Figure 6B). To examine whether these hair follicle-like structures were real hair follicles and whether these new hair follicles were regenerated from myeloid cells which were derived from splenocytes of GFP mice, we stained wounded skin with CD45 or GFP. The result shown in Supplement Figure S3 and Figure S4 demonstrated that many cells from hair follicle-like structures of healed wounds were either CD45-positive or GFP positive, suggesting myeloid cells participate in hair follicle regeneration. We then performed double immunofluorescent staining with antibodies of GFP and K14. As shown in Figure 7A, hair follicle-like structures shown in H & E stain (Figure 6B wound 1) were positive for both GFP and K14, further suggesting that GFP-positive splenocyte-derived myeloid cells were involved in new hair follicle regeneration. Moreover, isolated hair follicles from wounded skin of mice by collagenase were also double positive for GFP and K14 (Figure 7B). These data demonstrated that myeloid cells were able to regenerate hair follicles in wounded skin of mice.

**Figure 6.**
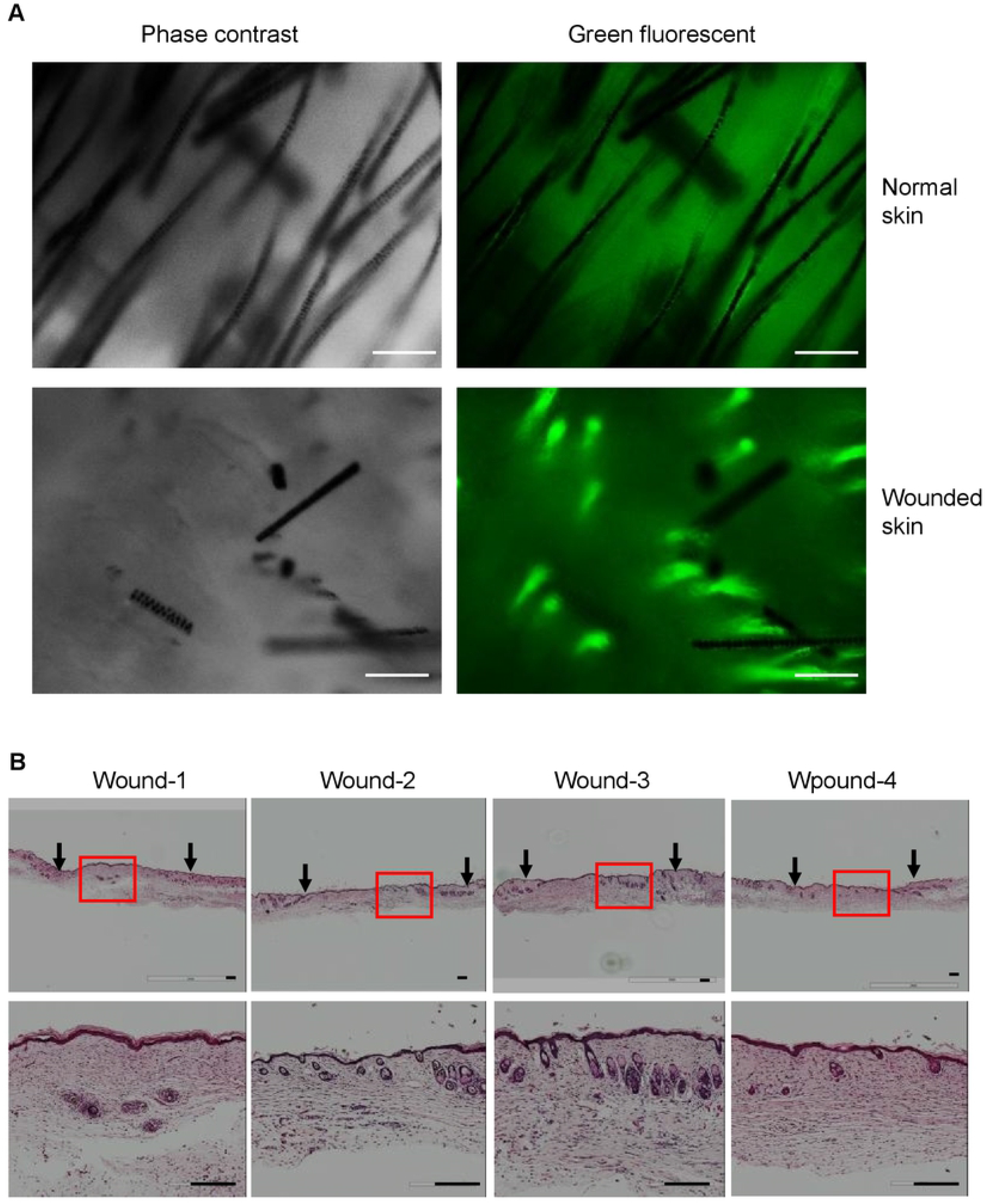
Hair follicle-like structures are revealed in wounded skin of mice received GFP-expressing myeloid cells for 4 weeks. Eight mm punch biopsy of excisional wounds were created in C57BL/6 mice which received one million GFP-expressed splenocyte-derived myeloid cells simultaneously. Four weeks after wounding, mice were euthanized, wounded skin were collected for examination of new hair follicle regeneration. (**A**) Images were taken under the fluorescent microscope in normal and wounded skin of the same mice received one million cells for 4 weeks. (**B**) H & E staining in wounded skin of mice received one million cells for 4 weeks. Arrows indicate wounding area. Top panels, magnification of 20 ×; Low panels, magnification of 100 ×. Scale bars, 100 μm.

**Figure 7.**
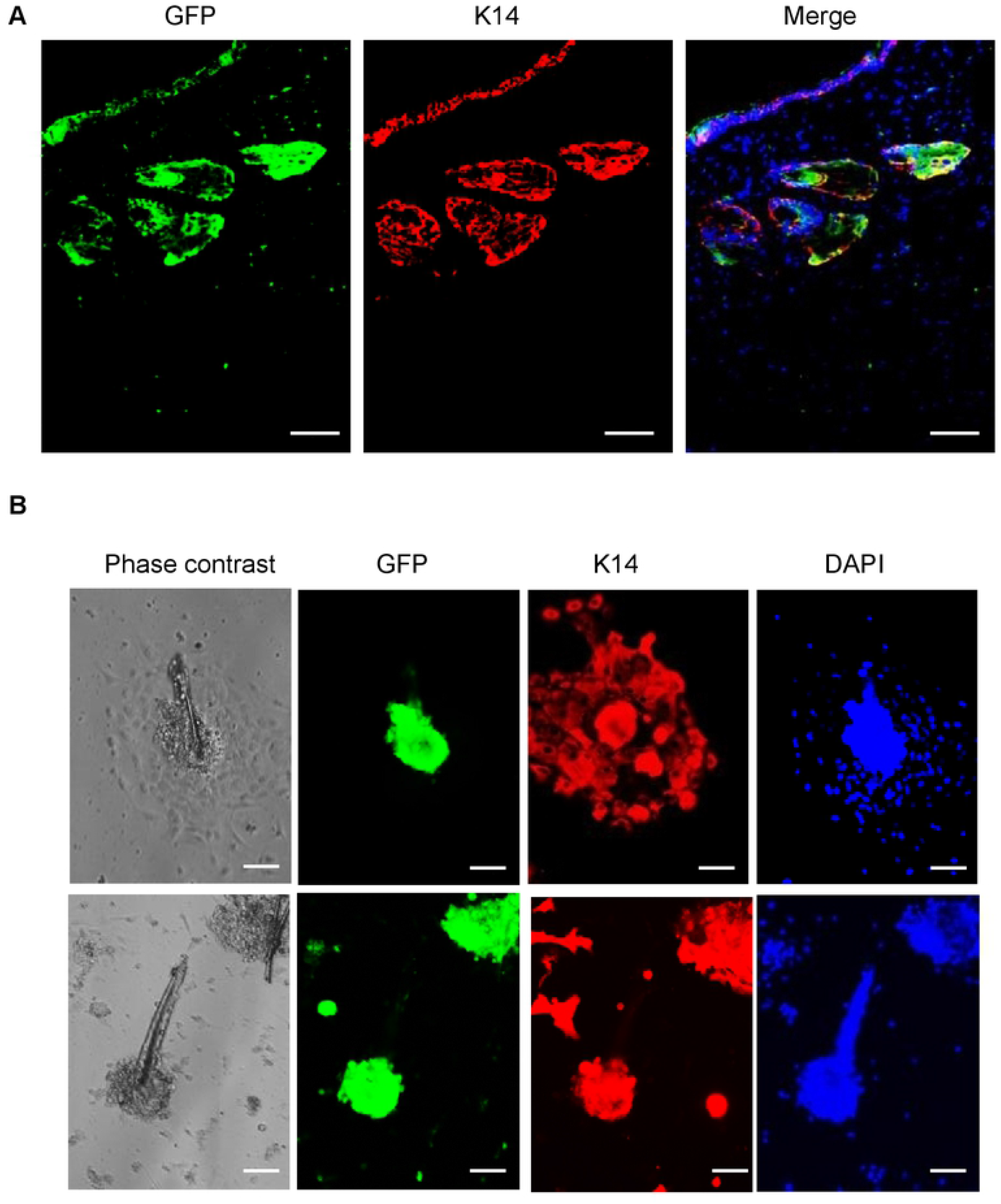
Detection of GFP-expressing myeloid cells in new forming hair follicle in wounded skin of mice. (**A**) Double immunofluorescent staining with keratin-14 (K14) (red) and GFP (green) antibodies in wounded skin of mice which received one million GFP-expressing splenocyte-derived myeloid cells for 4 weeks. (**B**) Double immunofluorescent staining with keratin-14 (K14) (red) and GFP (green) antibodies in cultured hair follicles isolated from the edge of wounds in mice received one million GFP-expressing myeloid cells for 4 weeks. DAPI (blue) was used as a nuclear counterstain. Scale bars in all images was 50 μm.

## DISCUSSION

In this study, by using double immunofluorescent staining with antibodies for hematopoietic cells, myeloid cells and differentiated keratinocytes as well as cell tracing approach, we have demonstrated that myeloid cells able to convert to skin keratinocytes and participate in hair follicle regeneration. Our data further support previous observations that tissue injury could induce hematopoietic cells to transdifferentiate into solid organ-specific cells [4–9].

It has long been known that myeloid cells such as neutrophils, monocytes and macrophages can be recruited to the wound sites and play a critical role in wound healing [19–21]. Wounds in mice expressing the human diphtheria toxin (DT) receptor under the control of the CD11b promoter and treated with DT to deplete CD11b-positive monocytes/macrophages, result in stalled healing [22]. Similarly, when mice expressing DT receptor under the control of a myeloid cell-specific lysozyme M promoter are wounded and depleted myeloid cells in different phase of healing by DT treatment, the granulation formation and epithelialization are delayed or stop healing [23]. Further, experimental evidence show that application of autologous monocytes/macrophages can significantly improve wound healing through the stimulation of angiogenesis and re-epithelialization in diabetic wounds in rats [24]. The mechanisms that myeloid cells improve wound healing are thought to be related to their scavenger ability, cytokines, chemokines, proteases and growth factors, released from these myeloid cells. However, here, our data strongly suggest a new mechanism of these myeloid cells in wound healing, in which myeloid cells may also improve wound healing *via* cell conversion.

The capacity of myeloid cells converting into endothelial cells, white adipocytes, osteoclasts and fibroblasts *in vivo* and *in vitro* has been documented [4,25–28]. Here we found that myeloid cells could also convert to keratinocytes, under the condition of skin injury. It is understandable that cells such as endothelial cells, adipocytes and fibroblasts can be converted from myeloid cells as they are from the same germ layer, mesoderm. Surprisingly, here we found that myeloid cells could also convert to keratinocytes. Myeloid cells need to cross the germline barrier to convert into keratinocytes which are from the ectoderm. The possible explanation for this type of conversion is dedifferentiation of myeloid cells into multipotent stem cells before conversion to keratinocytes. Indeed, we and other have previously demonstrated that hematopoietic cells could be dedifferentiated into stage specific embryonic antigen (SSEA)-1 and −3 positive multipotent stem cells when hematopoietic cells are cultured in a medium containing M-CSF [29–31]. We also reported that hematopoietic cell-derived SSEA-1 and SSEA-3 positive multipotent stem cells are transiently present at skin wound beds of mice [2]. After injury, recruited myeloid cells in the wound beds can be induced to be dedifferentiation by M-CSF released from injury-stimulating proliferating skin cells. We have also showed that topically application of M-CSF or its antibody in skin wounds can increase or delay healing process [2]. Therefore, it is likely that infiltrated myeloid cells are dedifferentiated into SSEA-positive multipotent stem cells induced by M-CSF and then myeloid cell-derived multipotent stem cells in wound beds further differentiated into skin fibroblasts, keratinocytes and other types of cells to contribute wound repair and skin appendage regeneration.

Hair follicles are thought to form only during development [32]. Loss of hair follicles are considered permanent. However, studies have shown that the hair follicles in some types of adult mice can be efficiently regenerated after skin injury [16, 17]. The types of cells to regenerate new hair follicles are still unclear. One study suggested that cells from the outside of hair follicle stem cells may be the cellular origin of new regenerated hair follicles [15]. Macrophages have been indicated to promote hair follicle regeneration [18]. In a recent study, using CD11b-DT receptor transgenic mice, Rahman and colleagues reported that wound-induced hair follicle regeneration is dependent on CD11b^+^F4/80^+^ cells [33]. Here, we provide further evidence to support that CD11b-positive myeloid cells are the cellular origin of converting keratinocytes and involved in wound-induced hair follicle regeneration. We have noticed that CD45-posiitve cells are dominant in new forming hair follicles. Furthermore, by injection of GFP-labeled splenocyte-derived myeloid cells to skin injury mice, we clearly showed that GFP-positive cells are dominant in new forming hair follicles. Taken together, our data strongly support that myeloid cells are at least one type of cells contributing to hair follicle regeneration after skin injury.

In summary, this study provides a new mechanism regarding to skin wound healing. We demonstrate that myeloid cells are involved in skin tissue repair and appendage regeneration through their conversion to skin keratinocytes. Our data also provide further proof to support a previous controversial topic, injury-induced hematopoietic stem cell transdifferentiation. The information obtained from this study not only clarifies the fundamental cellular and molecular mechanism of tissue repair and regeneration, may also provide new therapeutic options for treatments for chronic healing wounds *via* myeloid cell conversion.

## ACKNOWLEDGEMENTS

This study was supported by the Canadian Institutes of Health Researches (Funding Reference Number: 134214 and 136945). We thank Mr. Darrel Trendall for his kindly providing us euthanized C57BL/6-Tg (UBC-GFP) mice.

